# STRmie-HD enables interruption-aware HTT repeat genotyping and somatic mosaicism profiling across sequencing platforms

**DOI:** 10.64898/2026.03.21.713334

**Authors:** Alessandro Napoli, Niccolò Liorni, Tommaso Biagini, Agnese Giovannetti, Alessia Squitieri, Luca Miele, Andrea Urbani, Viviana Caputo, Antonio Gasbarrini, Ferdinando Squitieri, Tommaso Mazza

## Abstract

Short tandem repeat expansions in exon 1 of the *HTT* gene drive Huntington’s disease (HD) pathogenesis, with disease onset and progression heavily influenced by somatic mosaicism and sequence interruptions. While sequencing technologies enable repeat sizing, many computational tools lack the resolution to capture subtle interruption motifs and allele-specific somatic variation. We present STRmie-HD, an alignment-free, *de novo* framework for interruption-aware genotyping and quantitative profiling of somatic mosaicism at single-read resolution. The tool parses individual reads to quantify uninterrupted CAG tract length, CCG repeat content, and critical interruption variants, including Loss of Interruption (LOI) and Duplication of Interruption (DOI). Validated across Illumina, PacBio SMRT, and Oxford Nanopore platforms, STRmie-HD demonstrates high concordance with reference genotypes and superior sensitivity in identifying rare interruption patterns that conventional tools often overlook. Furthermore, it implements somatic mosaicism metrics to characterize repeat dynamics, successfully distinguishing the higher somatic expansion burden in brain tissues compared to peripheral blood. STRmie-HD offers a comprehensive and extensible solution for high-resolution molecular characterization of *HTT* variation, providing a robust framework for patient stratification and genetic research in HD.

**Graphical Abstract:** Graphical Abstract:
STRmie-HD flowchart. STRmie-HD is a comprehensive analytical framework that processes sequencing reads to analyze CAG/CCG trinucleotide repeats, interruption variants, and somatic mosaicism in the *HTT* gene. The workflow begins with sequencing reads (FASTA/FASTQ) that can undergo optional custom processing eq]based on the sequencing design. These reads are then fed into a regular expression-based engine (STRmie-HD) to identify CAG and CCG motifs. The identified motifs lead to the estimation of CAG/CCG alleles, visualized as distinct peaks representing different allele sizes, interruption variant assessment, and somatic mosaicism quantification. STRmie-HD produces an HTML output that wraps this information into a report.

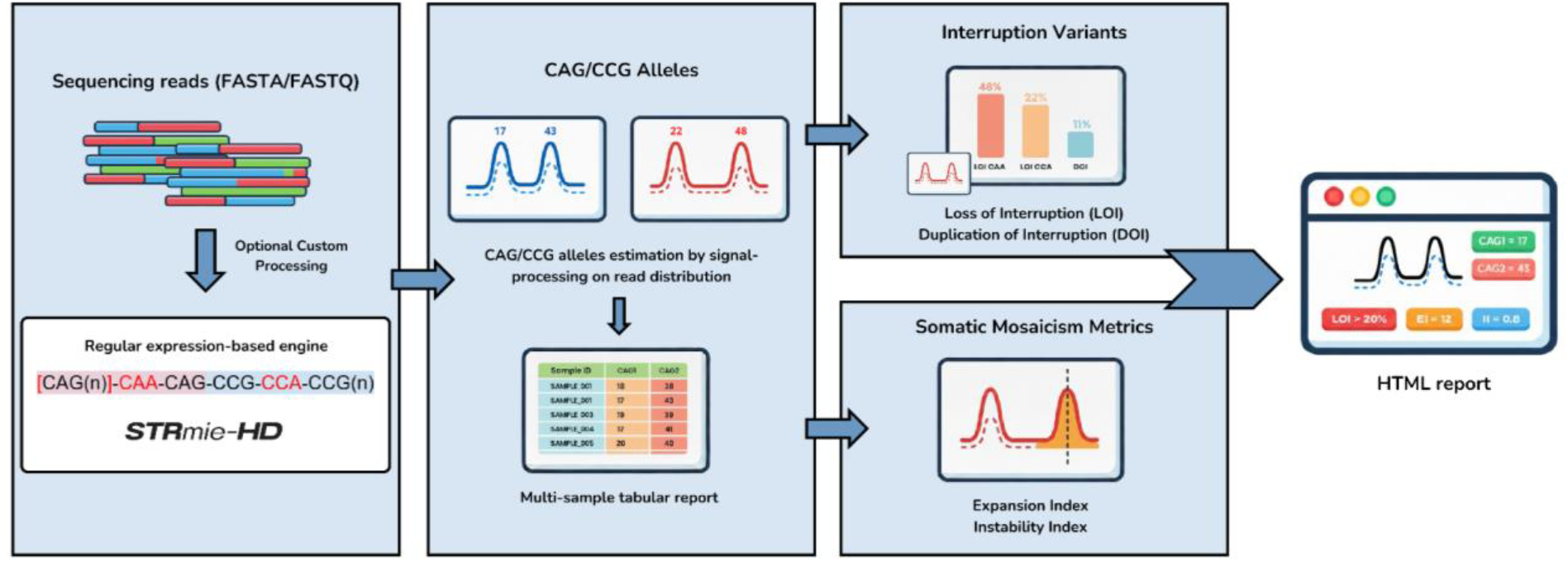

## Introduction

Huntington’s disease (HD) (OMIM #143100) is a rare autosomal dominant neurodegenerative disorder caused by an aberrantly expanded cytosine-adenine-guanine (CAG) trinucleotide tandem repeat [1], [2], [3] [4] in exon 1 of the *HTT* gene [5], located on chromosome 4p16.3. The *HTT*-exon 1 locus is composed of polymorphic CAG repeats encoding a glutamine-rich domain (*polyQ* tract) and of polymorphic CCG repeats encoding a proline-rich domain (*PRD* tract) in Huntingtin protein. The CAG repeat stretch is typically interrupted by a CAA triplet and followed by a final CAG repeat [6]. The CCG repeat stretch is typically interrupted by a CCA triplet and a variable series of CCG repeats encoding proline (P), which may stabilize the glutamine-rich tract [7]. Huntingtin is a multifunctional protein whose canonical function remains unclear [8], [9]. Mutated Huntingtin (mHTT) exhibits toxic properties that affect several key cellular and brain development processes, resulting in neurodegeneration and HD progression [10], [11].

HD is one of the best-studied repeat expansion disorders and affects an estimated 5–10 individuals per 100,000 globally [12], [13], [14], with approximately 6,000 cases in Italy [12], [15]. Approximately 41,000 people have symptomatic HD, and more than 200,000 are at risk of inheriting the disorder in the United States (Huntington’s Disease Society of America, 2023 https://hdsa.org/what-is-hd/overview-of-huntingtons-disease/). HD leads to progressive motor, cognitive, and psychiatric impairments, typically manifesting around the age of 45, although onset can range from childhood to later life. The length of the CAG repeat expansion strongly correlates with age at onset and disease severity: ≥40 repeats cause fully penetrant adult-onset HD, while 36–39 repeats show reduced penetrance. Juvenile-onset HD (JOHD), accounting for ∼5-10% of cases, often involves >60 repeats and presents with earlier and more severe symptoms, such as epilepsy and rapid decline [16], [17].

Although the inherited CAG length explains 42–71% of the age-at-onset (AOO) variance, the disease trajectory is not entirely predictable from the repeat length alone [18]. Somatic mosaicism, a tissue-specific variation in CAG repeat size, plays a critical role in HD [4]. The *HTT* gene exhibits both germline and somatic instability, with greater expansion observed when it is inherited paternally [19], [20], [21], [22]. This results in an earlier AOO and leads to a clinical feature known as *anticipation*, where the longer CAG repeat inherited by the progeny causes the disease phenotype to manifest much earlier in the offspring than in the carrier father [23]. Furthermore, pathological CAG triplets in the *HTT* gene vary in size among different cells within the affected tissues. For example, in HD brains, particularly in the cortex and striatum, CAG tracts can somatically expand into hundreds or thousands of units [24], [25], [26]. Mouse and human studies support this, with evidence suggesting that somatic CAG length better predicts age at onset than does germline size [27].

Other important modifiers are interruption variants (**Figure 1A**), especially the Loss of Interruption (LOI) variant [28], which is a single-nucleotide A>G change that converts the common CAACAG motif following the CAG repeat into a pure CAG tract. This variant does not alter polyglutamine (polyQ) length but significantly accelerates disease onset by up to 25 years, especially in reduced penetrance alleles (36–39 CAGs). LOI is enriched in symptomatic individuals within this range and segregates as a dominant trait, increasing somatic instability and underscoring that uninterrupted CAG length, not *polyQ* length itself, is a more accurate predictor of disease onset. A second LOI can also occur in the adjacent CCG tract. A distinct variant in this region that results in a longer interrupting sequence through the insertion of a second CAA-CAG motif [that is, (CAA-CAG)2], called DOI (Duplication of Interruption), has also been identified and linked to a delayed AOO in HD [7], [29], [30], [31]. This emphasizes the importance of detailed cis-structure characterization in genetic counseling and the development of predictive models.

**Figure 1:**
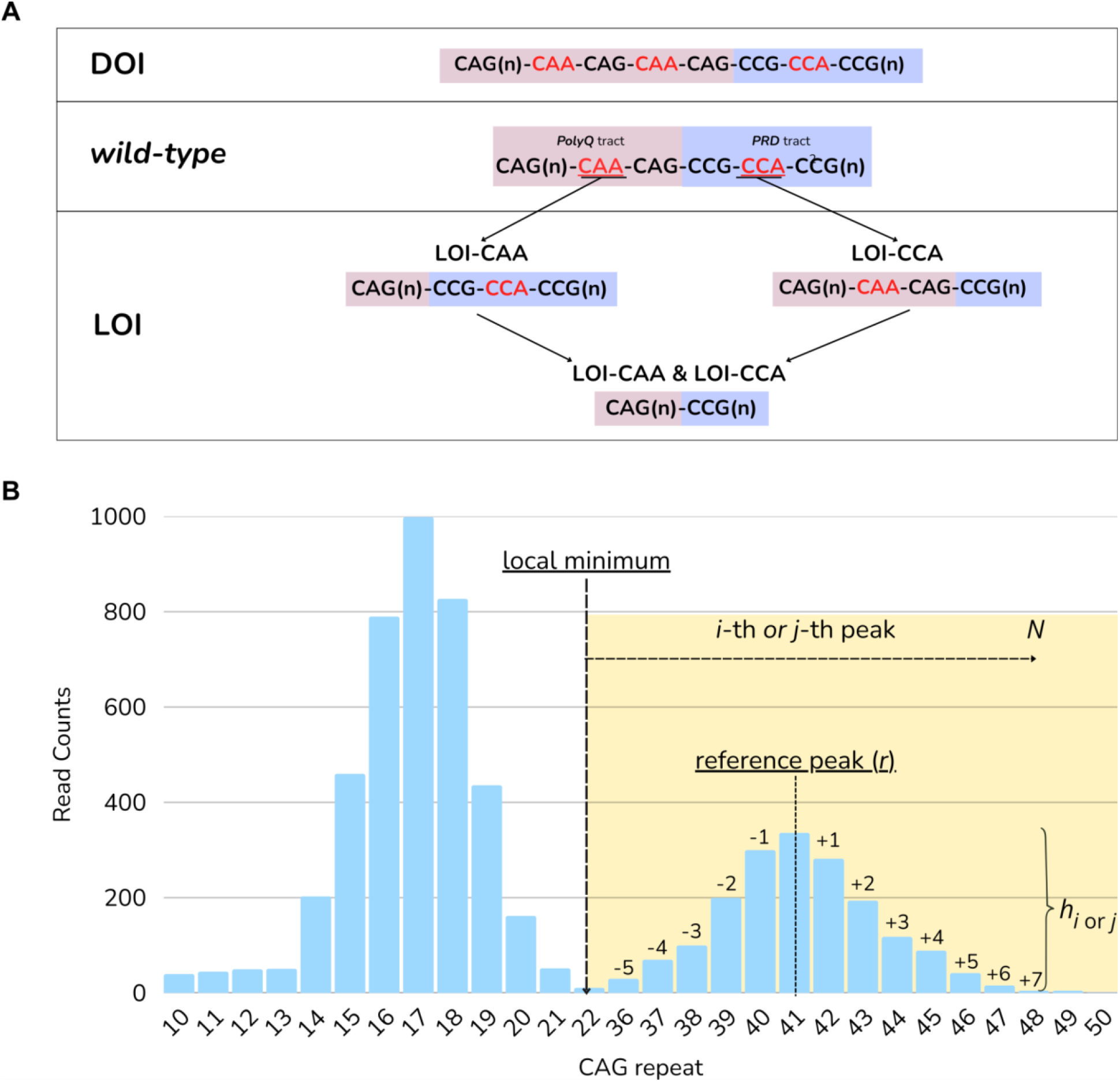
*HTT*-exon 1 repeat architecture and histogram-based indices: A) Exon 1 CAG–CCG tract and interruption patterns. B) Representative CAG-length histogram illustrating the local minimum between allelic modes, the reference peak at the expanded allele (*r*), per-bin height (*h*_*i*_), and the number of peaks (*N*) located to the right of the local minimum; these elements underlie the computation of the *Instability Index* and *EI*.

Diagnosis of HD relies on clinical evaluation, family history, and genetic testing. Conventional testing involves PCR amplification of *HTT* exon *1*, followed by fragment analysis or Sanger sequencing to determine the repeat length [32]. Triplet-primed PCR assays coupled with capillary electrophoresis (e.g., AmplideX PCR/CE *HTT* Kit) have simplified detection [33], [34], [35] and are now the gold standard in diagnostic settings. Small-pool or single-molecule PCR methods utilize serial dilutions and minimal DNA input to effectively detect rare, low-frequency expanded alleles from somatic mosaicism; however these techniques are notoriously labor-intensive and prone to contamination [36], [37]. More recently, Next-Generation Sequencing (NGS) has enabled more precise and high-throughput characterization of pathogenic expansions at a lower cost. Despite these advances, several challenges remain to be addressed. Standard short-read sequencing often fails to span the entire repeat tract, requiring indirect estimation from partially covering reads, as repetitive regions frequently exceed the typical read length and are difficult to align uniquely, representing an inherent limitation in the analysis of triplet-repeat expansion disorders, such as HD [38]. Even long-read whole-genome sequencing lacks sufficient depth and precision for expanded-repeat analysis. Thus, targeted sequencing remains the most reliable method for measuring repeat sizes and assessing mosaicism. Sequencing approaches such as Illumina MiSeq [39], Oxford Nanopore Technologies (ONT) [40] and Pacific Biosciences (PacBio) single-molecule, real-time sequencing (SMRT) [41], have enabled targeted high-resolution repeat analyses.

For instance, amplification-free targeted sequencing protocols using CRISPR/Cas9 on PacBio SMRT platforms have successfully resolved *HTT* repeat size and somatic mosaicism without PCR bias [42]. While sequencing triplet repeats is increasingly popular, bioinformatic analysis, especially for HD, remains a critical bottleneck, lacking accurate, consensus-driven computational strategies, and a comprehensive benchmarking of tools for triplet repeat characterization is still lacking [43], [44].

As outlined in a recent review by Maestri et al. [44], computational strategies for *HTT* CAG repeat profiling can be organized according to the sequencing design and input data, and by whether they implement *reference-based* (alignment-dependent) or *de novo* (reference-free) approaches. Within *reference-based* frameworks, widely used tools leverage locus-specific mapping, read-pair/fragment information, and coverage patterns to provide diploid genotypes; these include tools like ExpansionHunter [45], GangSTR [46], and TredPARSE [47] for Illumina WGS, ScaleHD [48] for Illumina MiSeq amplicons, and TRGT [49] or NanoRepeat [50] for high-accuracy long reads. Specialized reference-based workflows also include ONT methods such as DeepRepeat [51], NanoSTR [52] and NanoSatellite [53]. However, reliance on a reference can reduce sensitivity to highly expanded or interrupted molecules that map poorly, which is particularly relevant when somatic mosaicism generates a “long right tail” of rare alleles [44].

*De novo* approaches instead extract repeat information directly from read sequences without requiring genome-wide alignment (e.g., RepeatDetector [48], HMMSTR [49], RepeatHMM [54], CharONT2 [55]), which is advantageous for extreme expansions and characterizing repeat composition when alignment becomes unstable. Importantly, these tools can be further stratified by their support for (i) interruption variant detection, (ii) quantitative somatic mosaicism assessment from single-read length distributions versus only aggregate genotypes, and (iii) practical accessibility and consistency (installation/maintenance, documentation, reproducibility), all of which impact tool selection and benchmarking across datasets. Therefore, the suitability of any tool is not absolute but is contingent on the type of sequencing data, experimental strategy, and biological questions being addressed. Studies focusing on somatic instability or therapeutic monitoring benefit from tools that operate at single-read resolution and preserve allelic proportions, whereas diagnostic settings may prioritize robustness, throughput, and compatibility with clinical standards. As no single approach is universally optimal, method selection must align with both the technical nature and required resolution of the input data. Despite considerable advances, no single computational framework currently provides accurate HTT CAG/CCG genotyping together with quantitative characterization of somatic mosaicism and interruption variants [44]. In this study, we present STRmie-HD, a Short Tandem Repeat, alignment-free, regular expression-based mapping and identification engine that adopts a *de novo* approach to address the need for precise and comprehensive computational genotyping specifically designed for HD, with disease-tailored features such as interruption variant identification and numerical indices related to somatic mosaicism that are currently not fully supported by other computational tools.

## Materials and Methods

STRmie-HD is an analytical framework for the detection and quantification of CAG/CCG trinucleotide repeat expansions from targeted sequencing NGS data, with a specific focus on the *HTT*-exon 1 locus. For each input read, STRmie-HD parses the nucleotide sequence using a custom regular-expression scheme designed to capture the characteristic nucleotide architecture of the HTT exon 1 repeat tract (**Figure 1A**). It parses each read to extract the length of the uninterrupted CAG tract, the number of consecutive CCG repeats, and the presence or absence of specific interruption variants: LOI-CAA (loss of the canonical CAA interruption in the *polyQ* tract), LOI-CCA (loss of the typical CCA motif in the *PRD* tract), and DOI (duplication of CAACAG motifs in the *polyQ* tract). These annotations are compiled into a per-read dataframe, which serves as the basis for downstream individual and cohort-level summaries. At the sample level, STRmie-HD calculates the proportion of reads exhibiting interruption variant features and somatic mosaicism metrics, i.e., the *Expansion Index* (EI) and *Instability Index* (II).

For ONT data, STRmie-HD operates in a region-of-interest (ROI) parsing mode (enabled with the *--nanopore* flag) in which two locus-specific flanking anchor sequences are located within each read upstream and downstream the ROI tract. In this mode, STRmie-HD allows a user-defined maximum number of substitutions and indels during flank localization. This strategy mitigates the higher per-base error rates typically observed in ONT reads compared with short-read platforms. The sequence segment between the two anchors is then extracted and used for repeat assessment, optionally discarding reads whose extracted ROI does not meet a user-defined minimum fraction of in-frame CAG triplets (default 70%).

### Allele Detection

The identification of the two alleles within each sample is based on the distribution of CAG or CCG repeat lengths observed across the sequencing reads. This distribution is derived by counting the number of reads corresponding to each CAG/CCG repeat count, producing a histogram representing the allelic repeat-length profile for that sample.

STRmie-HD implements multiple strategies for automatic peak detection, allowing users to select the approach that is best suited to their datasets and experimental designs. By default, the two most prominent peaks are detected directly from the CAG/CCG repeat-count histogram using the *scipy*.*signal*.*find_peaks* Python function. Peaks are filtered based on a minimum distance criterion to ensure sufficient separation, and the two highest peaks, corresponding to the most frequent allele lengths, are selected and reported in ascending order.

Additionally, STRmie-HD provides alternative peak-detection strategies, including one based on a user-configurable cutpoint that bisects the repeat-count histogram into a “wild-type” and a “potentially expanded” zone and identifies local peaks within these newly defined regions. Here, we consider CAG cutpoints that are consistent with established clinical categories for *HTT*, i.e., ≤26 normal; 27–35 intermediate; 36–39 reduced penetrance; ≥40 full penetrance. This choice reflects a biologically motivated separation supported by clinical guidelines [56] but can be modified by the user.

As a final alternative for automatic peak detection, STRmie-HD provides a continuous wavelet transform (CWT) approach implemented using the *scipy*.*signal*.*find_peaks_cwt* module.

In all cases, STRmie-HD generates a report including histograms of CAG repeat distributions for each sample, which serve as the primary reference for result interpretation. By default, allele calls are derived automatically from the peak indices detected by the implemented algorithms. Given the importance of accurate peak identification for repeat-length determination, the generated histograms are provided to enable routine visual quality assessment of the detected peaks. While the automated procedures are sufficient for the vast majority of samples, this inspection step allows users to verify peak assignments in the presence of atypical distributions or borderline configurations. When necessary, STRmie-HD also provides functions for manual allele refinement and recalculation.

### Computation of Instability Index and Expansion Index

To quantitatively assess repeat instability and somatic expansion within each sample, STRmie-HD computes two principal metrics: the Instability Index (*II*) and the Expansion Index (*EI*). These indices are conceptually adapted from methodologies described in previous studies [57] [58] and are derived from the distribution of CAG repeat counts generated for each sample. Both metrics depend on the preliminary identification of the two representative CAG repeat lengths corresponding to the major alleles detected in the sample. Once these allelic peaks are established, STRmie-HD constructs a normalized histogram of the CAG repeat distribution and calculates the summary metrics from this distribution.

*II* quantifies the asymmetry of the CAG repeat distribution around the expanded allele and is calculated as follows (**Figure 1B**):

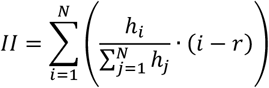

where *N* denotes the number of peaks located to the right of the local minimum between the two allelic modes and retaining all CAG bins to the right of that minimum, and *r* denotes the CAG repeat count of the expanded allele, defined here as the longer allelic mode. The *h*_*i*_ value represents the signal intensity (i.e., peak height) corresponding to the CAG repeat length *i*, while 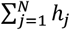 denotes the total signal across all *N* peaks. The term 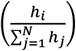 normalizes the height of each peak to the total height, thereby yielding the fractional contribution of each peak to the overall signal. This normalized value is then multiplied by (*i* − *r*), which quantifies the relative position of each peak with respect to the reference peak. Peaks to the right of the reference peak contribute positively (expansion), whereas those to the left contribute negatively (contraction).

The *EI* provides a quantitative measure of the accumulation of the CAG signal beyond the expanded allele, capturing the extent of downstream expansion and, therefore, the degree of somatic mosaicism. It is calculated as follows:

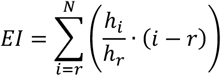

where *N* denotes the highest observed CAG repeat length among all the reads. h_r_ is the height of the reference peak at CAG length *r*. The term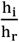 normalizes the peak height relative to the reference, thereby allowing a comparison across different signal intensities.

The sum is across all peaks with *i* ≥ *r* peaks to the right of the main peak. Thus, the summation yields a weighted measure of repeat expansion beyond the reference peak, integrating both the relative signal intensity and the extent of expansion.

### Benchmark datasets for CAG repeat size estimation

To evaluate the performance of STRmie-HD in correctly characterizing the *HTT*-exon 1 tract, we employed four distinct sequencing datasets. The first dataset (**Dataset 1**) consisted of 410 samples obtained from patients with HD, generated using Illumina MiSeq with a targeted paired-end approach, specifically designed to capture the *HTT* locus. This dataset included 399 blood samples and a small subset of cortex (n=6) and striatum (n=5) samples. Blood samples were provided by the LIRH Foundation collection, brain samples from the Leiden University Medical Centre [16]. For the benchmark, we restricted our analysis to 368 blood-derived samples for which an orthogonal experimental reference was available, namely PCR/CE-assessed CAG repeat sizes. Notably, a subset of seven samples of this Dataset featured independent orthogonal validation for the loss of the canonical CAA interruption (LOI-CAA), providing a critical ground-truth set to evaluate STRmie-HD’s precision in resolving these complex and clinically significant interruption variants. The second dataset (**Dataset 2**) consisted of 22 PacBio SMRT-sequenced lymphoblastoid cell lines (GEO accession number: GSE199005) previously analyzed in the RepeatDetector study, originally obtained from the Coriell Institute for Medical Research cell repository (https://www.coriell.org/) [59]. For this dataset, RepeatDetector-based allele reconstructions were considered the concordance baseline as the independent orthogonal genotyping was not available for all samples on the Coriell Institute for Medical Research cell repository (either not available or only available upon request) and the dataset was originally introduced to benchmark repeat sizing on PacBio SMRT sequencing data in the RepeatDetector original publication. The third dataset (**Dataset 3**) consisted of 10 synthetic samples designed and generated to emulate a spectrum of clinically relevant scenarios in a naïve single-end, long-read design. These synthetic sequences were specifically engineered with predefined LOI and DOI configurations to rigorously benchmark the engine’s ability to not only identify CAG repeat size but to also provide precise, read-level quantitative estimates of interruption mosaicism across a controlled range of repeat architectures. The fourth dataset (**Dataset 4**) consisted of 11 ONT samples (SRA accession: PRJNA678742) of HD cell lines obtained from the Coriell Institute for Medical Research cell repository previously characterized in the DeepRepeat original paper, for which a reference Sanger sequencing ground-truth was available [51]. Sanger sequencing genotypes used as reference were obtained from the original publication, specifically from Supplementary Tables ST3 and ST4, and were used as the ground truth for allele size comparison [60].

#### Ethics approval

Study protocols and informed consent forms for patients and healthy controls were approved by the Institutional Review Board of the LIRH Foundation on 28th October 2022 (protocol no. 10.281022) and on February 7, 2023 (protocol no. 12.070223). The study was conducted in accordance with the Declaration of Helsinki.

### *HTT*-targeted Next Generation Sequencing protocol for Dataset 1 and bioinformatics processing

All **Dataset 1** samples were sequenced using an ultra-high-depth sequencing protocol run on an Illumina MiSeq platform, specifically targeting the *HTT* gene locus. This approach involved multiple reads targeting different sections of the repeat tract to accurately estimate the number of CAG repeats. In particular, the *HTT*-exon 1, including the repeated tract, was amplified from genomic DNA samples and sequenced using the Illumina paired-end MiSeq platform. Primer sequences used to amplify the *HTT* region were: HTT_1444_F ATGAAGGCCTTCGAGTCCC HTT_1444_R GGGCTGAGGAAGCTGAGGA.

For all reactions, 1X KAPA HiFi HotStart ReadyMix (Roche Diagnostics) containing 0.5 U per 20 μL reaction buffer, dNTPs (0.3 mM of each dNTP at 1X), MgCl2 (2.5 mM at 1X), and stabilizers with 6% DMSO addition. The thermal protocol was composed by first 18 cycles 95°C 30s, 61°C 30s (decreasing 0.5°C per cycle), 72°C; and then 22 cycles 95°C 30s, 52°C 30s, 72°C 30s and final extension 72°C 5 min.

The paired-end reads generated by this targeted NGS protocol were merged prior to downstream analysis using PEAR (Paired-End reAd mergeR, v0.9.11) [61]. This preprocessing step was required because STRmie-HD and RepeatDetector operate on single continuous read sequences spanning the repeat tract. Merging the forward (R1) and reverse (R2) reads therefore allowed reconstruction of the original DNA fragment into a single sequence covering the full CAG repeat region, avoiding the need to analyze paired reads independently and subsequently reconcile separate repeat-length estimates. PEAR identifies statistically supported overlaps between paired reads by integrating nucleotide matches, sequencing error probabilities, and base quality scores. The resulting merged reads spanned the CAG repeat tract and were used as direct input for STRmie-HD and RepeatDetector to estimate repeat lengths at the single-molecule level. PEAR was run with default parameters optimized for Illumina data, enforcing a minimum overlap of 10 bp and requiring statistically significant overlap support (p-value ≤ 0.05). Reads that did not meet these criteria, often due to insufficient overlap in very large expansions or sequencing artifacts, were excluded from the downstream analysis. Summary statistics of the PEAR merging performance across samples are reported in **Supplementary Table 1**. Tools capable of directly handling paired-end reads were executed using their native input requirements.

### Dataset 3 synthetic generation

A synthetic dataset of samples containing 10,000 sequencing reads, was generated to simulate the repeat architecture of the *HTT* gene. Each read was designed to represent exon 1 of *HTT*, incorporating variable-length CAG and CCG repeat tracts, as well as a specific Loss of Interruption (LOI) pattern at the junction between these repeats. CAG repeat lengths were sampled from two normal distributions centered around typical allele sizes, by default 17 and 49 repeats, with a default standard deviation of 2. To introduce allelic diversity, an additional subset of repeat lengths was randomly selected between 10 and 100. All CAG values were clipped to fall within the range of 1–100 repeats. CCG repeat lengths were independently assigned and drawn from two predefined values (default: 7 and 10 repeats). We modeled six LOI configurations that reflected known and hypothetical interruptions between the CAG and CCG tracts. These included canonical and variant motifs, such as CAACAGCCGCCA, CAGCAGCCGCCG, CAGCAGCCGCCA, CAACAGCCGCCG, CAACAGCAACAGCCGCCA, and CAACAGCAACAGCCGCCG. The relative abundance of each LOI type was controlled using user-defined percentages, which were set to 30%, 20%, 15%, 10%, 20%, and 5%, respectively. For each read, the final sequence was constructed by concatenating the CAG repeat, selected LOI motif, CCG repeat, and terminal CCT sequence. The reads were either padded or truncated to a fixed length of 300 nucleotides and assigned a constant Phred quality score to mimic high-quality sequencing data. The complete set of reads was shuffled randomly and saved in FASTQ format, simulating realistic sequencing variability while preserving the control over the repeat structure and composition. This synthetic dataset enabled precise benchmarking of repeat sizing, LOI detection, and overall tool accuracy across a wide range of biologically relevant scenarios.

In particular, we generated 10 samples, of which one was labeled as “healthy” (15/25), two as “intermediate” (12/34, 16/30), two as “reduced penetrance” (17/37, 15/39), two as “full penetrance” (19/40, 23/49), two as “high expansion” (17/74, 18/84) and one as “very high expansion” (18/102). Of these 10 samples, two were generated as DOI samples, three as LOI samples, and one as an LOI-CCA sample, while the remaining samples were generated without reads carrying interruption variants.

### Tools selection criteria for comparative benchmarking

The benchmarking tools selected to evaluate STRmie-HD were chosen based on their compatibility with the specific sequencing input data available for each dataset. The selection process followed the criteria outlined by Maestri et al. [44], namely: (1) the required type of input data; (2) the methodological framework, distinguishing between *de novo* and *reference-based* approaches; and (3) the level of output granularity (single-read *versus* allele-level). In addition to these criteria, we introduced further inclusion parameters: (4) relative obsolescence, excluding tools that have been superseded by more advanced algorithms according to recent literature; (5) support for the detection of interruption variants; (6) capability to detect and/or quantify somatic mosaicism; (7) support for diploid genotyping; (8) availability of appropriate benchmarking dataset; and (9) the current maintenance status of the software. Consistent with this selection strategy, we decided to include one *reference-based* and one *de novo* comparator for each dataset.

To assess the profiling accuracy of STRmie-HD on **Dataset 1**, we compared its performance with ScaleHD (*reference-based*, optimized for Illumina MiSeq data) and RepeatDetector (*de novo*). For the PacBio SMRT sequencing data in **Dataset 2**, TRGT was selected as the *reference-based* comparator, alongside RepeatDetector as the *de novo* method. The analysis of **Dataset 3** was limited to RepeatDetector as the sole comparator, due to the synthetic single-end long-read design of the dataset, which was incompatible with conventional *reference-based* approaches. Finally, for the ONT sequencing data in **Dataset 4**, benchmarking was conducted against DeepRepeat as the *reference-based* method and CharONT2 as the *de novo* alternative. The extensive reasons for tool selection are outlined in **Supplementary Table 2**.

### Input Data Processing and Software Configuration

For **Dataset 1**, STRmie-HD and RepeatDetector (Linux version 1.0.15eb445) were executed on PEAR-processed single-end FASTQ/FASTA reads, whereas ScaleHD (v.1.0) analyzed the original paired-end FASTQ files due to its mandatory input files requirements with the parameters *atypical_realignment, genotype_prediction*, and *snp_calling* enabled, producing a tabular report containing genotype predictions (**Supplementary Material**). For **Dataset 2**, BAM files for TRGT (v4.0.0) were generated by aligning FASTA reads to the hg38 human reference genome using minimap2 (ver 2.3) (preset *map-hifi*, retaining secondary alignments).

TRGT was run with the *HTT* repeat BED file (chr4:3074876-3074933, ID=HTT;MOTIFS=CAG,CCG;STRUC=<HTT>) and produced allele predictions in VCF format (**Supplementary Material**). STRmie-HD and RepeatDetector processed the FASTA reads directly. For **Dataset 3**, both STRmie-HD and RepeatDetector were run on the synthetic FASTA files directly. For **Dataset 4**, DeepRepeat predictions were retrieved directly from the original publication [51], while STRmie-HD was executed using its dedicated *--nanopore* flag to account for the higher base-calling error rates characteristic of ONT sequencing with default parameters. CharONT2 (ver 1.1.0) was run to cluster ONT reads at the STR locus into two dominant haplotype groups and generate one consensus sequence per group (allele 1 and allele 2). For each allele-consensus, CAG repeat length was obtained from the CharONT2 allele report by selecting the triplet-repeat call (period size = 3) with consensus motif CAG and using its reported copy number as the allele size; repeat calling was performed under a stringent configuration (e.g., TRF parameters *2,500,500, 80, 10, 50, 500*) to prioritize precise quantification of the core CAG tract. The two haplotype consensus groups were reported as the two alleles for each sample. For **Dataset 1, Dataset 2** and **Dataset 3**, RepeatDetector was executed in both *restrictive* mode, aimed at precise CAG repeat quantification, and *permissive* mode, intended to enhance detection of interruption variants. For each sample, a CAG-repeat-length histogram was generated, and the two highest peaks were interpreted as the primary alleles. Because the *permissive* mode indirectly identifies interruptions by reporting characteristic shifts in repeat counts, for interruption variants detection we used the *restrictive* profile prediction as the baseline count (designated as *n*) and we then interpreted the *permissive* mode’s output distribution relative to this baseline: a peak at *n*+1 repetitions was considered a LOI allele, a peak at *n*+5 repetitions a DOI allele, and a peak at *n*+3 repetitions a canonical allele. In all datasets, for both STRmie-HD and RepeatDetector, the CAG-repeat-length histogram was considered as the main output, and the two highest peaks were interpreted as the two main alleles.

### Benchmark metrics

The quantitative assessment of prediction accuracy was performed by calculating error metrics between predicted and reference repeat sizes. For the 368 blood-derived samples in **Dataset 1** with ground truth genotypes, performance was evaluated using both Mean Absolute Error (MAE) and Root Mean Square Error (RMSE). While MAE reflects the average prediction error, RMSE provides a more stringent measure by penalizing larger, clinically significant deviations. 95% confidence intervals (CIs) for MAE and RMSE were derived via non-parametric bootstrapping (1,000 iterations), defined by the 2.5th and 97.5th percentiles of the resampled error distributions. For this large cohort, we also computed Spearman’s rank correlation coefficient (ρ) to evaluate the strength of the monotonic association between predicted and ground-truth values and assessed the ability of each method to correctly classify alleles into four clinically relevant phenotypic categories (normal, intermediate, reduced penetrance, and full penetrance) using macro-averaged precision and recall scores. In contrast, for the smaller cohorts (**Datasets 2** and **4**, with *n*=22 and *n*=11 respectively), accuracy was assessed using MAE alone. We intentionally omitted RMSE, Spearman correlation, and categorical classification for these datasets due to the low sample size. All computations were performed in Python (v3.13) using a combination of the *pandas, numpy, scikit-learn*, and *scipy* libraries.

### Tissue-level comparisons of somatic mosaicism indices

To investigate the biological relevance of *II* and *EI*, we performed a comparative analysis across the three available tissue types in all 410 **Dataset 1** samples: blood (n=399), cortex (n=6), and striatum (n=5). Because somatic mosaicism is a known hallmark of HD, with brain tissues exhibiting greater repeat expansion than peripheral tissues, this comparison served as a critical biological contextualization of our metrics. Boxplots were generated for each index to visualize the distribution of values within each tissue group. Given the non-normal distribution of the data and the small sample sizes for the brain tissues, we used the non-parametric Kruskal-Wallis test for statistical comparisons. We conducted a post-hoc pairwise comparison to assess the significant differences between groups: blood versus cortex, blood versus striatum, cortex versus striatum, and blood versus a consolidated “brain” group (cortex and striatum combined). This analysis allowed us to explore whether the indices could capture the expected tissue-specific patterns of somatic instability in patients with HD.

## Results

### STRmie-HD is highly accurate in estimating CAG repeat size in Illumina MiSeq HD samples

To evaluate the performance of STRmie-HD on clinically relevant targeted Illumina MiSeq samples (**Dataset 1**), we compared the computed versus PCR/CE-assessed CAG repeat lengths for both alleles (CAG1, unexpanded and CAG2, expanded) using RMSE and MAE. STRmie-HD consistently performed equally or better than the other tools across these error metrics in **Dataset 1** (**Supplementary Table 3, Dataset 1**). It computed a median CAG repeat size of 17 for the unexpanded alleles and a median CAG repeat size of 43 for the expanded alleles across all 368 benchmarked samples (**Figure 2B)**. Along with RepeatDetector, it achieved low RMSE and MAE (**Figure 2A; Table 1**) for both CAG1 and CAG2. In contrast, ScaleHD exhibited an uneven performance, with reduced accuracy in detecting larger CAG repeats. We also evaluated the accuracy of the tools in quantifying CAG repeat lengths across two alleles by computing Spearman’s rank correlation coefficients with the PCR/CE-assessed repeat sizes (**Figure 2C**). STRmie-HD showed high correlation overall, achieving ρ = 0.93 (p-value = 2.54e-168) for CAG1 and ρ = 0.93 (p-value =1.81e-162) for CAG2, together with the *restrictive* RepeatDetector profile, which also performed well (CAG1: ρ = 0.93, p-value = 2.54e-168; CAG2: ρ = 0.92, p-value = 8.09e-159), although it was slightly less consistent for expanded alleles. ScaleHD showed good performance for CAG1 (ρ = 0.90, p-value = 7.69e-138) but performed poorly for CAG2 (ρ = 0.1, p-value = 5.9e-02).

**Figure 2.**
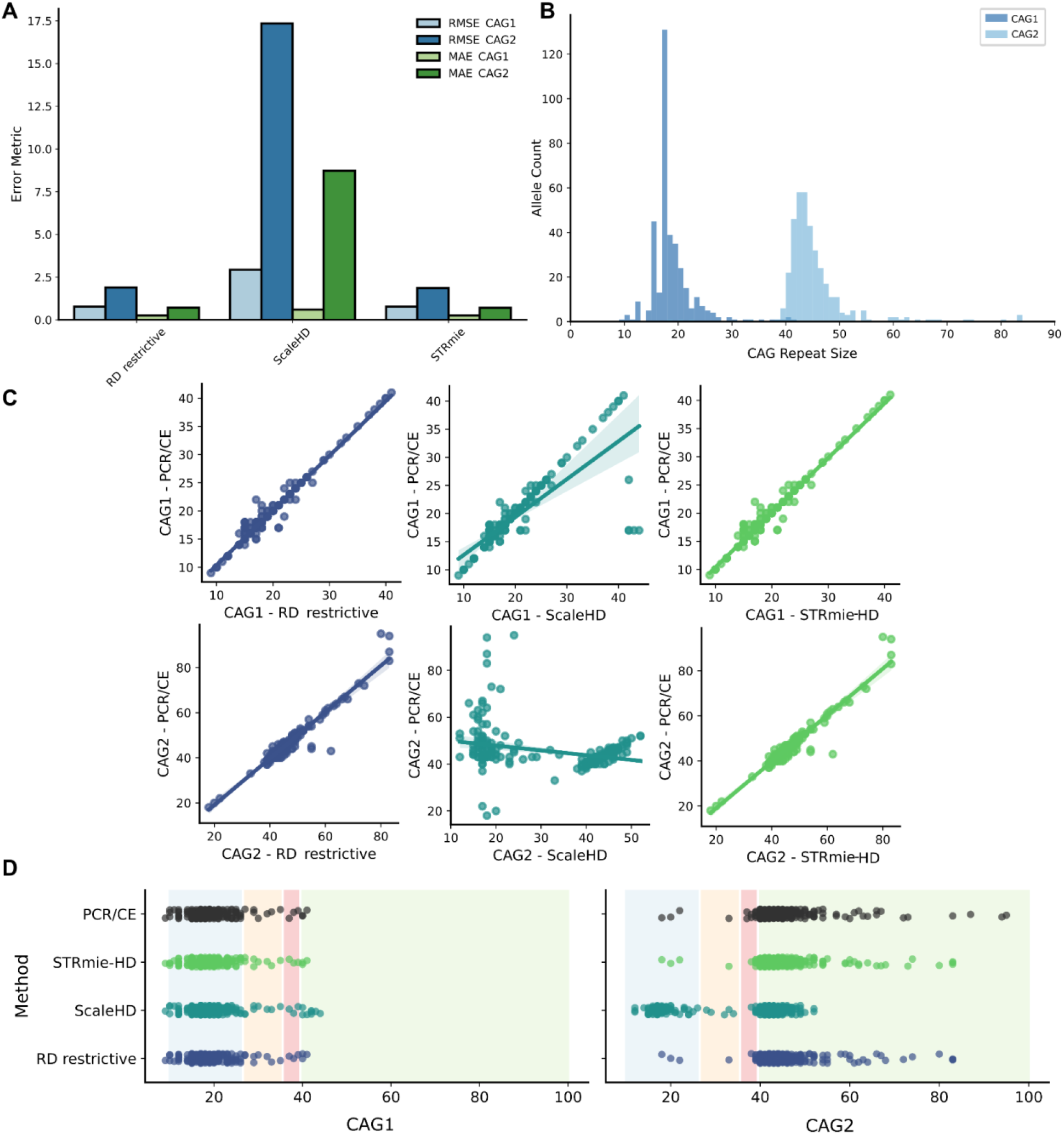
Benchmarking of *HTT* CAG repeat sizing tools on Dataset 1. A) Bar plot comparing the performance of STRmie-HD against RepeatDetector (RD) and ScaleHD using Root Mean Square Error (RMSE) and Mean Absolute Error (MAE) for both the unexpanded (CAG1) and expanded (CAG2) alleles. B) Histogram showing the distribution of allele sizes predicted by STRmie-HD across all 368 benchmarked samples in **Dataset 1**. C) Scatter plots of CAG repeat sizes predicted by each tool versus the ground-truth PCR/CE-assessed genotypes. The diagonal line represents a perfect correlation, and the shaded area surrounding each regression line represents the 95% confidence interval for the regression prediction. D) Allele distribution plots comparing the categorization of CAG repeats by each tool into clinically relevant ranges: normal, intermediate (light grey line), reduced penetrance (grey line), and full penetrance (black line). Each dot represents an allele from a single sample, providing a visual assessment of the classification accuracy against ground-truth PCR/CE ranges.

**Table 1:** Comparison of genotyping accuracy for different methods. The table shows the Root Mean Square Error (RMSE) and Mean Absolute Error (MAE) for the STRmie-HD, RepeatDetector in restrictive mode and ScaleHD methods on the expanded and unexpanded alleles (CAG1 and CAG2). Values are presented with 95% confidence intervals (CI).

| Metric | STRmie-HD | RD restrictive | ScaleHD |
| --- | --- | --- | --- |
| RMSE CAG1 | 0.77 (95% CI: 0.6–0.93) | 0.77 (95% CI: 0.62–0.91) | 2.86 (95% CI: 1.52–4.07) |
| RMSE CAG2 | 1.83 (95% CI: 1.21–2.48) | 1.87 (95% CI: 1.26–2.52) | 17.35 (95% CI: 15.32–19.49) |
| MAE CAG1 | 0.26 (95% CI: 0.20–0.34) | 0.26 (95% CI: 0.19–0.33) | 0.60 (95% CI: 0.33–0.94) |
| MAE CAG2 | 0.70 (95% CI: 0.54–0.90) | 0.72 (95% CI: 0.55–0.91) | 8.75 (95% CI: 7.30–10.26) |

Next, we evaluated the accuracy of each method in categorizing *HTT* CAG repeat sizes into clinically relevant phenotypic categories assessed with the ground truth as a direct consequence of their CAG repeat length predictions (**Figure 2D**). Based on this classification approach, STRmie-HD and the *restrictive* profile of RepeatDetector achieved the best overall performance. STRmie-HD achieved precision scores of 0.97 (CAG1) and 0.96 (CAG2), with corresponding recall values of 0.99 and 0.89, respectively. RepeatDetector in *restrictive* mode showed identical performance, with high precision (0.97 for CAG1, 0.96 for CAG2) and solid recall (0.799 and 0.815) (**Figure 2D**). ScaleHD delivered excellent performance for CAG1 (precision: 0.997; recall: 0.99) but lacked precision for CAG2 (precision: 0.48; recall: 0.80). Collectively, these results support STRmie-HD as an accurate and robust solution for genotyping the *HTT* CAG repeat region when adopting this sequencing design, with RepeatDetector following closely.

Finally, leveraging STRmie-HD’s capability to quantify CCG content in addition to CAG repeats, we analyzed the CCG allelic distribution in all 410 **Dataset 1** samples (**Supplementary Table 4, Dataset 1**). We found the CCG allelic distribution to be very limited and dominated by CCG=7, with the most common genotypes being 7/7 (60.2%; 247/410), 7/10 (29.5%; 121/410), and 10/10 (3.4%; 14/410), with all other combinations accounting for ∼6.9%.

### STRmie-HD precisely reconstructs the CAG repeat profiles of SMRT-sequenced and synthetic samples

To evaluate performance on PacBio SMRT-sequencing data, we analyzed **Dataset 2**. STRmie-HD accurately reconstructed the allelic CAG repeat distributions for all 22 samples and showed strong concordance with previously reported RepeatDetector profiles generated under *restrictive* settings. Specifically, STRmie-HD achieved a combined MAE of 0.16, with a perfect match for the unexpanded allele (MAE = 0.00) and high precision for the expanded allele (MAE = 0.32) compared to the RepeatDetector in *restrictive* profile’s predictions (**Figure 3A, B, Supplementary Table 3, Dataset 2**).

**Figure 3.**
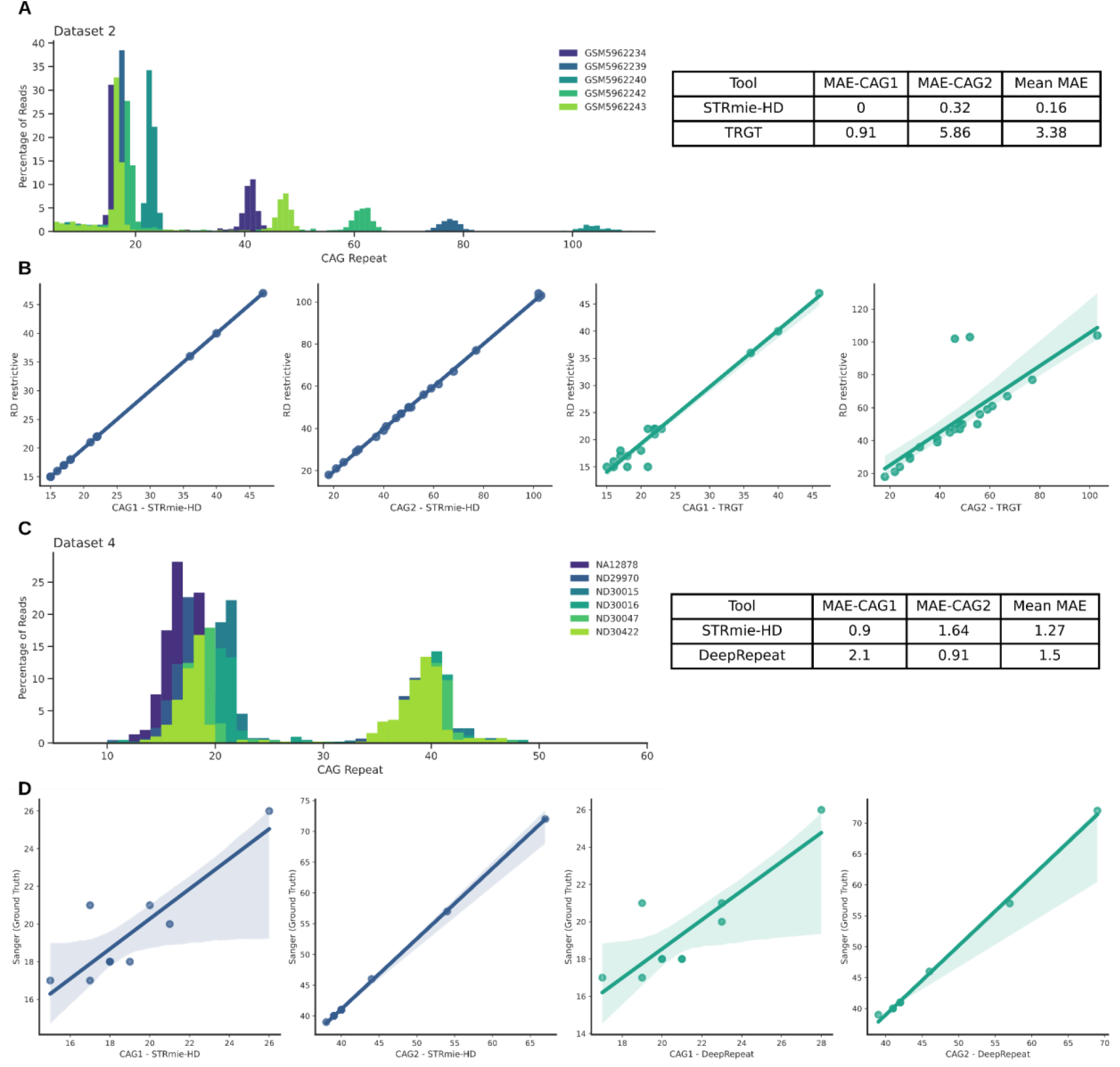
Reconstruction of CAG profiles by STRmie-HD and RepeatDetector (*restrictive*) on Dataset 2 and Dataset 4: Histogram plot representing the CAG profiles of **Dataset 2 (A)** and **Dataset 4 (C)** samples obtained with STRmie-HD. Only a selection of five samples per dataset is shown for clarity. Scatter plots of CAG of **Dataset 2 (B)** and of **Dataset 4 (D)** repeat sizes predicted by each tool versus the ground-truth genotypes (Repeat Detector predictions for **Dataset 2** and Sanger for **Dataset 4**).

Furthermore, while TRGT also showed strong concordance for the unexpanded alleles (MAE = 0.91), we noted a significant difference in its quantification of the expanded alleles. This was reflected in a substantially higher MAE of 5.86 for Allele 2. For example, TRGT estimated an allele being 46 CAGs for the sample GM13506, whereas both STRmie-HD and RepeatDetector counted 102 repeats. Similarly, TRGT counted 52 CAGs for an allele of sample NA20245, in stark contrast to the 103 repeats identified by the other two methods. Similar results were obtained on a different dataset of 11 PacBio SMRT-sequenced samples [42], where STRmie-HD recovered the exact CAG repeat size determined with fragment analysis (**Supplementary Table 5**). Considering only the samples for which a ground-truth genotype was available in the Coriell repository (10/22), RepeatDetector in *restrictive* mode returned an MAE of 0.3 for the unexpanded allele and 0.4 for the expanded allele. STRmie-HD showed comparable accuracy for the unexpanded allele (MAE = 0.3) and a slightly higher MAE for the expanded allele (MAE = 0.7). In contrast, TRGT exhibited larger deviations, with an MAE of 1.3 for the unexpanded allele and 1.5 for the expanded allele. Two samples were excluded from this specific MAE calculations: GM14044, owing to its exceptionally long second allele reported in the Coriell repository (750 CAG), as highlighted by the RepeatDetector authors, and GM13506, where both RepeatDetector and STRmie-HD consistently detected 102 repeats, a sharp contrast to the 48 repeats documented by Coriell. We also analyzed the CCG allelic distribution in all 22 **Dataset 2** samples (**Supplementary Table 4, Dataset 2**). The distribution was highly constrained and dominated by CCG=7; the most common genotypes were 7/7 (63.6%; 14/22) and 7/10 (27.3%; 6/22), with all other combinations accounting for 9.1%.

Finally, we evaluated the allele-calling performance of STRmie-HD on synthetic samples (**Dataset 3**). STRmie-HD correctly recovered the CAG repeat counts embedded in the sequences, matching the performance of RepeatDetector under its *restrictive* profile across all tested scenarios (**Supplementary Table 3, Dataset 3**).

Collectively, these results indicate that STRmie-HD reproduces established RepeatDetector-based allele profiles on PacBio SMRT sequencing data while maintaining comparable or improved accuracy relative to alternative methods in the subset of samples for which an external ground-truth genotype is available.

### STRmie-HD maintains reliable CAG repeat calling in Oxford Nanopore data

To assess the performance of STRmie-HD on noisier long-read sequencing platforms, we evaluated its ability to accurately reconstruct *HTT* CAG repeat alleles from targeted ONT data. Using **Dataset 4** samples, we compared CAG repeat size estimates generated by STRmie-HD, DeepRepeat and CharONT2 against Sanger-based CAG sizing reported in the DeepRepeat study [51]. Overall, STRmie-HD showed high concordance with the ground truth, yielding an average MAE of 1.27, performing comparably to DeepRepeat (MAE of 1.50). Performance nuances were observed between the two tools: STRmie-HD demonstrated higher accuracy for the unexpanded allele (MAE = 0.90 vs. 2.10 for DeepRepeat), whereas DeepRepeat was more precise in sizing the expanded allele (MAE = 0.91 vs. 1.64 for STRmie-HD) **(Figure 3C, D, Supplementary Table 3, Dataset 4**).

CharONT2 showed limited completeness for diploid CAG reconstruction on ONT data. Across the 11 samples (**Supplementary Table 3, Dataset 4**), CharONT2 returned both allele CAG genotypes in only 3/11 cases (ND30626, ND30015, ND33947). In these samples, the unexpanded allele was generally close to the Sanger-based genotype (absolute deviations of 2–3 repeats), while the expanded allele showed a wider spread (2–7 repeats), including an underestimation in ND33947 (33 vs 40 CAG). In additional 4/11 samples (ND31551, ND30422, ND30047, NA12878), CharONT2 returned a CAG call for only the unexpanded allele (ND31551: 20 vs 18 CAG; ND30422: 20 vs 18 CAG; ND30047: 21 vs 18 CAG), whereas the expanded allele was not recovered. No CAG allele was returned for 5/11 samples (ND33392, ND30016, ND29970, ND40534, GM04723).

Finally, leveraging STRmie-HD’s capability to quantify CCG content in addition to CAG repeats, we analyzed the CCG allelic distribution in all 12 samples (**Supplementary Table 4, Dataset 4**). The most common genotypes were 7/10 (58.3%; 7/12) and 7/7 (25.0%; 3/12), with less frequent genotypes collectively accounting for 16.7% of cases.

### STRmie-HD finely characterizes the interruption variant profile of synthetic and clinically relevant samples

Next, we evaluated the ability of STRmie-HD in quantifying the proportion of reads carrying interruption variants in **Dataset 1, Dataset 2** and **Dataset 3. Dataset 4** was excluded from this analysis as the inherent error profile of nanopore sequencing data limits its suitability for the robust detection of sequence interruptions. STRmie-HD identified 39 out of the 410 **Dataset 1** with a high proportion of reads containing LOI-CAA, LOI-CCA, or DOI variants (**Figure 4; Supplementary Table 6, Dataset 1**). We then compared these predictions with those of RepeatDetector (*permissive* mode) and ScaleHD. For comparison with RepeatDetector’s *permissive* mode, we used its representations of a canonical allele (*n*+3 CAG repetitions, with n being the number of CAG repetitions predicted by the RD’s restrictive profile), LOI-CAA allele (*n*+1 CAG repetitions), and DOI allele (*n*+5 CAG repetitions). For ScaleHD, we considered the “*Typical/Atypical*” flagging assessment for the presence of interruption variants in the data. However, neither RepeatDetector nor ScaleHD could identify the precise percentage of reads carrying a specific interruption variant. Among these 39 samples, ScaleHD flagged only 32 samples as having an *Atypical* allele, whereas RepeatDetector’s predictions and distributions consistently aligned with STRmie-HD, reporting a shift to *n*+1 for LOI-positive samples and to *n*+5 for DOI-positive samples, thereby confirming the presence of these variants. No shift was reported for LOI-CCA positive samples as the *PRD* tract is not processed by RepeatDetector. In several samples (e.g., HD4501–HD55601), STRmie-HD identified high LOI-CAA and LOI-CCA proportion of reads (∼30–35%); however, ScaleHD flagged these as *Typical*, possibly due to limited resolution in capturing partial interruption mosaicism.

**Figure 4.**
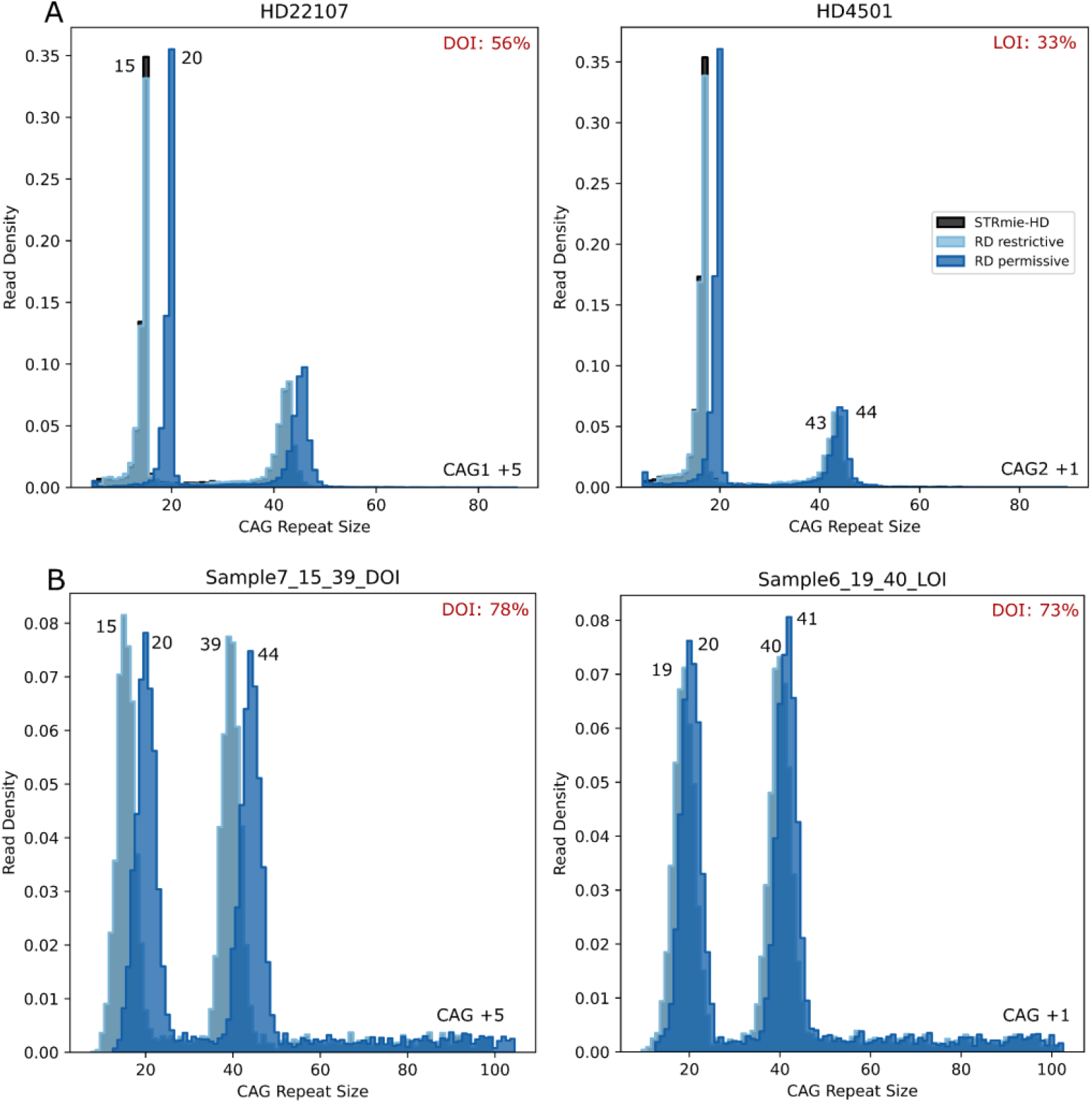
Characterization of interruption variants in Dataset 1 and Dataset 3 samples: Histograms distributions of two representative samples from **Dataset 1 (A)** and two representative samples from **Dataset 3 (B)** identified by STRmie-HD as carrying interruption variants. The STRmie-HD profiles (black, in the background) were compared to RepeatDetector’s (RD) *restrictive* (light blue) and *permissive* (blue) outputs, which indicate interruptions by characteristic shifts of +5 (DOI) or +1 (LOI). Interruption variants percentages identified by STRmie-HD are reported in red on the top right of each plot.

The accuracy of STRmie-HD’s interruption calls was definitively confirmed by a benchmark subset of seven samples (HD4501, HD4503, HD4504, HD47701, HD55601, HD73301, and HD75901) with prior orthogonal validation for LOI-CAA. While these samples were previously verified to lack the canonical CAA triplet, STRmie-HD successfully identified all seven as LOI-CAA-positive, providing precise read fractions between 30.9% and 48.5%. Notably, while RepeatDetector’s permissive mode corroborated these findings via an indirect +1 CAG shift, ScaleHD failed to detect any abnormality, misclassifying all seven as “*Typical*”. These results underscore STRmie-HD’s unique capability to quantitatively resolve validated interruption-loss events with single-read resolution where conventional tools falter.

In **Dataset 2**, STRmie-HD identified only one sample (GSM5962228) with a high proportion of reads carrying an interruption variant, particularly a LOI in the CCG tract (38,8%) (**Supplementary Table 6, Dataset 2**), which was not discernible with RepeatDetector. When applied to **Dataset 3**, STRmie-HD accurately recovered the ground-truth proportions of embedded LOI and DOI reads across all 10 generated samples (**Supplementary Table 6, Dataset 3**).

Overall, these results underscore one of the key strengths of STRmie-HD in providing not only precise identification of interruption variants but also quantitative estimates at the single-read level.

### The Expansion Index mirrors the degree of somatic mosaicism in blood, cortex and striatum samples

We used STRmie-HD to compute *II* and *EI* in **Dataset 1, 2** and **4** to assess the degree of somatic mosaicism across samples (**Supplementary Table 7**). In **Dataset 1**, *II* exhibited a broad range, spanning from a minimum of -8.76 to a maximum of 6.47 (**Figure 5A**). Negative values of this metric indicate a shift in the distribution of repeat lengths toward shorter fragments relative to the expanded allele peak, reflecting a bias toward contraction events within the population of somatic cells. Conversely, higher positive values suggest an increased representation of longer-than-expected repeats, indicating a bias toward somatic expansion. *EI* quantifies the relative abundance of secondary peaks that represent somatically expanded CAG alleles, reflecting the degree of somatic mosaicism, and ranges from 0.067 to 45.98 across all the analyzed samples (**Figure 5B**). Interestingly, the highest *EI* values were observed in samples derived from brain tissues, specifically the cortex and striatum (notably HD123st, HD123cx, HD255st, and HD255cx, as well as HD208cx, HD257cx, and HD257st). In **Dataset 2**, *II* ranged from a minimum value of -5.60 (GSM5962242) to a maximum value of 9.98 (GSM5962245). *EI* ranged from 0.58 (GSM5962228) to 54.10 (GSM5962245). In **Dataset 4**, *II* ranged from a minimum value of -1.60 (ND30626) to a maximum value of 0.50 (NA12878), whereas *EI* ranged from 1.0 (NA12878) to 68.8 (ND29970).

**Figure 5.**
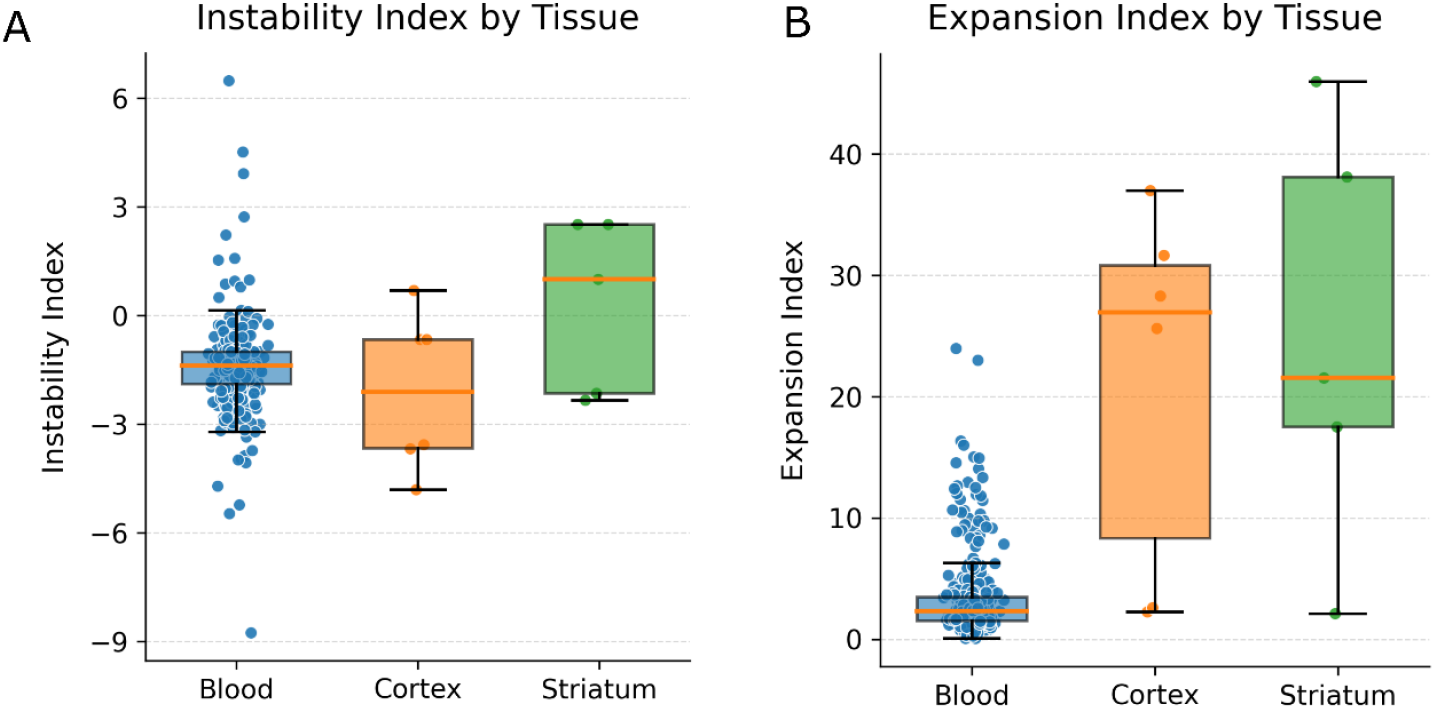
Tissue-specific differences in somatic mosaicism: **(A)** *Instability Index* and **(B)** *EI* measured in samples derived from blood, cortex, and striatum tissues. Boxplots display the distribution of each index across the sample types, with individual sample values overlaid as jittered dots.

To biologically contextualize this further, we compared the *EI* across samples derived from the cortex, striatum, and blood tissues from **Dataset 1** using the Kruskal–Wallis statistical test. Blood samples displayed significantly lower *EI* values than both the cortex (p-value =0.0032) and striatum (p-value =0.0028), and the difference remained highly significant even when blood samples were compared to the combined group of brain tissues (cortex + striatum; p-value =3.2e-05) (**Figure 5B**). The *EI* did not differ significantly between the cortex and striatum (p-value =0.8551). No statistically significant differences were observed in *II* between any of the sample groups: blood vs. cortex (p-value =0.7332), blood vs. striatum (p-value =0.2596), blood vs. brain (p-value =0.6141), and cortex vs. striatum (p-value =0.1003) (**Figure 5B**). These findings suggest that, in this dataset, the *EI* clearly distinguishes blood from brain tissues, reflecting a greater burden of somatic expansions in brain tissue relative to blood, whereas the *II* appears more consistent across tissues, with no strong evidence of tissue-specific differences in expansion-contraction bias.

## Discussion

The characterization of *HTT* exon *1* currently relies on a diverse range of sequencing inputs, from short-read Illumina MiSeq amplicons to high-resolution long-read data from PacBio SMRT sequencing and Oxford Nanopore. Because each platform presents a unique technical profile, ranging from the high-throughput robustness of short-read platforms to the single-molecule resolution required to track somatic instability in long-read data, the selection of the proper computational analytical framework has historically been dictated by the specific nature of the input sequencing technology. Despite these technological options, a standardized and versatile pipeline capable of bridging these distinct data formats to accurately resolve repeat size, interruptions, and mosaicism simultaneously has remained an unmet need [44].

In this study, we introduced STRmie-HD to address this gap, demonstrating consistent and robust performance across these diverse sequencing designs. Within the datasets and tool set assessed here, STRmie-HD achieved superior or comparable accuracy in quantifying PCR/CE-assessed genotypes in 368 Illumina MiSeq samples (**Dataset 1**) compared to the *reference-based* ScaleHD and the *de novo* RepeatDetector. Furthermore, STRmie-HD demonstrated high precision in reconstructing CAG profiles from 22 PacBio SMRT-sequencing samples (**Dataset 2**). In this context, it performed better than TRGT, a state-of-the-art *reference-based* method, particularly on expanded alleles, while matching the performance of RepeatDetector, the *de novo* tool that originally utilized this specific dataset to benchmark its own capabilities. These results were further validated through the analysis of 10 synthetic samples (**Dataset 3**), which simulate an ideal, error-free, single-end long-read design. In this controlled scenario, STRmie-HD achieved performance levels comparable to RepeatDetector, confirming its efficacy in handling high-resolution, long-read data architectures. Finally, it maintained reliable CAG calling on 11 ONT samples (**Dataset 4**), performing comparably to DeepRepeat and providing complete diploid outputs more consistently than CharONT2 in this dataset. Taken together, these results support STRmie-HD as a single framework that can be applied across multiple *HTT* targeted-sequencing designs, while acknowledging that performance depends on dataset characteristics and tool-specific input constraints.

A key feature of STRmie-HD is its ability to perform direct, quantitative, read-level characterization of interruption variants, such as Loss of Interruption (LOI) in both the *polyQ* and *PRD* tracts and Duplication of Interruption (DOI), which are critical modifiers of HD pathogenesis, addressing a profiling gap that is not fully met by existing bioinformatic tools. Unlike tools that infer interruptions indirectly, such as RepeatDetector, or ScaleHD, which provides a simple, non-quantitative classification, STRmie-HD reports interruption variants quantitatively at single-read level as the exact percentage of reads carrying LOI-CAA, LOI-CCA or DOI motifs. This substantially improves interpretability by enabling direct quantification of interruption variants at the single-read level. In particular, with ScaleHD only providing a binary “*Typical/Atypical*” output, without distinguishing the extent of the atypical pattern, and RepeatDetector requiring two independent runs (*restrictive* and *permissive* modes) followed by manual comparison of their shifted peaks to infer the presence of interruption variants, making interpretation dependent on user expertise and additional post-processing, STRmie-HD delivers an immediate, unambiguous, and interpretable quantification of interruption variants mosaicism. Inaccurate CAG length estimations and interruption variants identification can impact more than just age-of-onset predictions; they are critical for clinical trial eligibility, which is often based on specific CAG sizes or CAG–age product scores. Accordingly, interruption-aware genotyping may reduce ambiguity in research pipelines and, with appropriate validation, could support clinical research workflows in which repeat sizing and interruption status are relevant. Accurate LOI detection is vital because uninterrupted CAG tracts are linked to a worse phenotype, anticipated age of onset and increased somatic instability [28]. The advantage of STRmie-HD is evidenced by its performance on seven orthogonally validated samples lacking the canonical CAA interruption (LOI-CAA). While ScaleHD failed to flag these cases and RepeatDetector provided only indirect instability signatures, STRmie-HD accurately recovered all seven. Beyond binary detection, STRmie-HD offered high quantitative resolution, identifying these variants in 30.9% to 48.5% of the read pool. This single-read precision is crucial for characterizing the low-frequency mosaic variants that modulate Huntington’s disease progression.

In addition to CAG sizing and interruption profiling, STRmie-HD also enables direct joint genotyping of the adjacent CCG tract, thereby extending locus resolution across the full *HTT* exon *1* repeat architecture. More broadly, resolving both CAG and CCG components within the same analytical framework provides a more complete molecular description of *HTT* exon *1* and may facilitate the identification of clinically relevant sequence configurations involving interruption patterns across the combined CAG–CCG region.

Beyond genotyping, STRmie-HD implements two metrics, the EI and the II, to quantitatively describe the dynamics of somatic mosaicism directly from sequencing data. Although the *EI* and *II* were originally defined and characterized in previous literature [57], [58], to our knowledge no existing HD-oriented tool provides a fully documented and peer-reviewed implementation of the *EI* and *II* metrics. STRmie-HD therefore enables systematic application of both indices to high-depth targeted sequencing data. The *EI* quantifies the degree of rightward skewness in the expanded allele distribution, reflecting the extent of outlier expansions, whereas the *II* measures the asymmetry of the distribution around the expanded allele, capturing the dynamic range of somatic events. We investigated the biological relevance of these indices by analyzing samples from different tissues known to exhibit differential mosaicism [24], [25], [26]. In particular, the EI was significantly higher in brain tissues (cortex and striatum) than in blood. This finding aligns with the established knowledge that the central nervous system, particularly these regions, is a hotspot for somatic expansion in HD, thereby confirming the utility of these metrics in capturing tissue-specific disease processes and regional vulnerabilities.

In addition to its performance and key innovations, an important practical advantage of STRmie-HD is its ease of use. The tool requires no reference alignment and provides a simple command-line interface. The GitHub repository (https://github.com/mazzalab/STRmie-HD) includes detailed documentation, extensive examples, and ready-to-run workflows, making STRmie-HD accessible not only to bioinformaticians but also to laboratories with limited computational expertise.

Nevertheless, users should be aware that input preprocessing choices and sequencing platform error profiles can influence read-level summaries and allele distributions, and parameters may require dataset-specific tuning.

Although STRmie-HD was developed specifically for HD, its core framework is extensible. Its alignment-free methodology, which utilizes a customizable regular expression–based engine for parsing repeat tracts and interruption motifs, can be readily adapted for genotyping other tandem repeat disorders. Tandem repeat expansions have been implicated in over 60 human diseases, many of which have neurological symptoms. By modifying the core regular expression to target the unique repeat motif and flanking sequences of a different locus, the tool’s analytical pipeline may be repurposed to analyze expansions and interruptions in other genes, such as the CAG repeat in *ATXN1* (Spinocerebellar ataxia type 1, SCA1) [62], [63], the CGG repeat in *FMR1* (Fragile X syndrome) [64], the CTG repeat in *DMPK* (Myotonic Dystrophy Type 1) [65], [66], [67], or the complex GGGGCC hexanucleotide repeat in *C9orf72* (Amyotrophic Lateral Sclerosis and Frontotemporal Dementia) [68], [69]. Furthermore, many of these disorders also exhibit significant somatic mosaicism, a key feature that STRmie-HD is designed to quantify through its *II* and *EI*. By providing detailed characterization of repeat length, interruptions, and somatic variability, STRmie-HD may facilitate research studies, support patient stratification in clinical trials, and improve molecular characterization of repeat expansion disorders.

More broadly, STRmie-HD aligns with the need for computational methods that are not only accurate, but also practical, scalable, and reproducible for molecular research. Previous work has emphasized the importance of accessible computational recipes for genome research [70], cost-conscious high-performance computing strategies [71], and parallel approaches to manage the intrinsic complexity of biological data analysis [72]. In this context, STRmie-HD contributes a disease-focused, alignment-free framework that aims to combine methodological transparency with applicability across multiple sequencing platforms, supporting robust repeat profiling in Huntington’s disease research.

## Author Contributions

Alessandro Napoli: Formal analysis, Methodology, Software, Writing – original draft. Niccolò Liorni: Formal analysis, Methodology, Software, Writing – original draft. Tommaso Biagini: Supervision, Writing – review & editing. Agnese Giovannetti: Resources, Writing – review & editing. Alessia Squitieri: Resources, Investigation. Luca Miele: Investigation, Supervision. Andrea Urbani: Investigation, Supervision. Viviana Caputo: Supervision, Methodology, Writing – review & editing. Antonio Gasbarrini: Investigation, Supervision. Ferdinando Squitieri: Investigation, Supervision, Resources, Writing – review & editing. Tommaso Mazza: Conceptualization, Supervision, Software, Writing – original draft.

## Acknowledgments

We acknowledge the Italian Ministry of Health (RC2026). We thank Hannah S. Bakels, Susanne T. de Bot and Willeke M.C. van Roon-Mom for providing patients’ brain samples to FS. We are grateful to patients and families who take part in the research programs of Fondazione Lega Italiana Ricerca Huntington (LIRH).

## Data Availability

Documentation and tool releases are available at: https://github.com/mazzalab/STRmie-HD.

**Dataset 1** FASTQ files are available at SRA archive (**PRJNA1400480**).

**Dataset 2** FASTA files are available at the GEO database (**GSE199005**).

**Dataset 3** FASTQ files are available at Zenodo: https://doi.org/10.5281/zenodo.18346811.

**Dataset 4** FASTQ files are available at SRA archive (**PRJNA678742**).

